# Cross-View Latent Integration via Nonparametric Gamma Shrinkage Factor Analysis

**DOI:** 10.64898/2026.01.02.697340

**Authors:** Hassan Akell, Małgorzata Łazęcka, Dinesh Adhithya Haridoss, Miriam Urban, Eike Staub, Ewa Szczurek

## Abstract

Factor analysis is a dominant paradigm for multi-omic heterogeneous data, but is challenged by partially redundant signals and noise across views and by an unknown true number of factors. We present CLING, an unsupervised multi-view factor model with hierarchical Bayesian sparsity priors: a product-of-Gammas prior inducing cumulative column-wise shrinkage (increasing with factor index) coupled with a Gamma–Gamma local-precision hierarchy on loadings yielding heavy-tailed marginals. This pairing enables automatic factor selection by adaptively deactivating unsupported factors while retaining active ones during inference, and induces selective sparsity that allows salient loadings to escape shrinkage while collapsing negligible ones. As a fully conjugate hierarchical model, CLING admits a scalable variational inference algorithm for multi-view data. Across synthetic benchmarks and multiomics datasets, CLING recovers more accurate factors and more informative loadings while explaining at least as much variance as competitive multi-view baselines; on glioblastoma gene expression and DNA methylation data, CLING identifies pathways linked to tumor subtype and patient age. Source code: https://github.com/szczurek-lab/CLING.

## 1 Introduction

Large-scale studies in medicine and biology increasingly profile the same samples using multiple heterogeneous data *views*, such as gene mutations, gene expression, and DNA methylation. Integrating such multi-omics measurements is essential for understanding complex biological systems. Linear multi-view factor models provide a principled framework for explaining such data using a small number of hidden *factors, loadings* that link features to factors, and a *noise* term with feature- and view-specific precision.

Construction of adequate sparsity priors on the loadings is critical for multi-view factor analysis. Sparse and stable loadings are necessary to capture shared and view-specific variation with interpretable structure. Sufficient prior adaptivity is needed to account for imbalanced signal-to-noise ratios across views and features. Appropriate column-wise shrinkage of loading matrices aids selection of the unknown number of active factors *K*. An inadequate estimate of *K* obscures the recovered latent structure: underspecifying *K* compresses signal, whereas overspecifying allows surplus factors to absorb noise. Finally, the choice of sparsity priors dictates key mathematical properties, such as conjugacy, which directly affect the efficiency of inference algorithms. In particular, while variational inference offers scalable runtimes, Gibbs sampling approaches generally scale less favorably.

Sparsity priors proposed previously suffer from a number of limitations. Classical approaches such as MOFA (Argelaguet et al., 2018) promote sparsity through the introduction of hard zeros via spike–and–slab priors and apply non-cumulative, view-global shrinkage using automatic relevance determination (ARD). Hard zeros aid readability but can diminish weak-but-real loadings; non-cumulative shrinkage does not account for imbalanced signal-to-noise ratios and can obscure true signals. More recent MuVI (Qoku and Buettner, 2023) promotes sparsity via a softer regularized horseshoe prior, and often reports binary active feature sets by thresholding continuous loadings for interpretation; it also relies on non-conjugate black-box variational inference, adding optimization and implementation complexity. A promising alternative is soft shrinkage toward zero via the multiplicative gamma process (MGP), originally developed for single-view infinite factor models (Bhattacharya and Dunson, 2011), which imposes progressively stronger column-wise sparsity as the factor index *k* increases.

To address these challenges, we introduce CLING (Cross-view Latent Integration via Non-parametric Gamma Shrinkage), a multi-view factor model with hierarchical Bayesian sparsity priors. To control the overall scale of loading columns, we apply a product-of-Gammas prior, resulting in column-wise cumulative shrinkage. This construction gently switches off unsupported loading columns, retains supported ones, and calibrates each view’s effective rank, eliminating the need to hand-tune *K*. The shrinkers on elements of the loading matrices adapt sparsity at the feature–factor level. To this end, we place a simple Gamma–Gamma prior with heavy-tailed marginals (including half-*t* and half-Cauchy cases), which effectively shrinks small loadings strongly while leaving large, meaningful loadings essentially unchanged, yielding *selective sparsity* and robust, interpretable loadings. To accommodate heterogeneous noise, each feature in each view is assigned its own noise precision. Together, the local–global shrinkage and flexible precision provide complementary adaptation across columns, entries, and features. Consequently, the model respects view- and feature-level signal-to-noise differences and jointly explains variation across heterogeneous views. All components are chosen to preserve conjugacy (Gaussian–Gamma–Gamma–Gamma), yielding closed-form mean-field variational updates with monotone evidence lower bound (ELBO) improvement and an efficient variational inference algorithm.

Empirically, these modeling choices yield substantial practical benefits. In simulations with increasing numbers of factors, higher noise levels, and stronger sparsity, CLING more accurately recovers latent factors and loadings while explaining as much or more total variance than competing methods. On real multi-omics datasets, CLING produces factor-based sample representations that improve prediction of biologically and clinically relevant covariates. Applied to glioblastoma data (Brennan et al., 2013), CLING identifies gene expression and DNA methylation pathways associated with tumor subtype and patient age. Collectively, these results position CLING as a reliable default framework for integrative analysis of large, noisy, and heterogeneous multi-omics datasets.

## 2 Related works

Factor analysis (FA) offers a general framework for uncovering latent structure in data. Classical linear baselines such as PCA (Hotelling, 1933) and Tucker decomposition (Tucker, 1966) focus on low-rank reconstruction but fix the latent dimensionality *K a priori*. In contrast, Bayesian FA, particularly in its nonparametric form, employs sparsity-inducing priors that allow the data to determine *K*. A prominent approach places an Indian Buffet Process prior over a binary loading mask, as in Nonparametric Sparse Factor Analysis (NSFA) (Knowles and Ghahramani, 2011). A complementary line of work uses MGP shrinkage to define infinite factor models (Bhattacharya and Dunson, 2011; Durante, 2017), with extensions to Pitman-Yor process mixtures (Murphy et al., 2019) and to multi-study factor analysis (De Vito et al., 2021; Grabski et al., 2023; Hansen et al., 2025), where inference has been performed using both Gibbs sampling and variational inference. Additionally, heavy-tailed global–local shrinkage priors are widely used (Carvalho et al., 2009; Makalic and Schmidt, 2016).

However, many modern datasets are multi-view, with multiple modalities measured on the same samples, for which single-view FA is inadequate. Beyond MOFA and MuVI, newer models include FACTM (Łazęcka and Szczurek, 2025), which augments FA with a correlated topic component and supervised rotations to integrate structured and simple views; BSFP (Samorodnitsky et al., 2024), which performs joint multiview factorization with prediction; and JAFAR (Anceschi et al., 2024), which introduces a dependent cumulative shrinkage process prior to distinguish shared and view-specific factors.

## 3 Methods

### 3.1 Background

#### Multi-view FA

Given *M* views 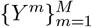 measured on the same *N* samples, probabilistic multi-view FA posits a shared latent representation *Z* ∈ ℝ^*N×K*^ and view-specific loadings 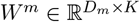, with

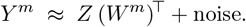

a formulation studied in Bayesian group and multi-block FA prior to its adoption in multi-omics (Virtanen et al., 2012; Klami et al., 2014) and later popularized in this context by MOFA (Argelaguet et al., 2018).

A standard route to structured sparsity combines spike-and-slab priors with ARD. For view *m*, feature *d*, and factor 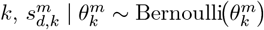,

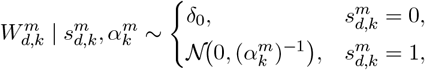

with 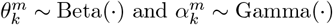. Tying the inclusion probability 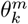 and slab precision 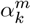 across *d* encourages entire columns of *W*^*m*^ to deactivate when unnecessary, yielding clear zero/nonzero structure and strong feature selection.

#### MGP shrinkage

In single-view Gaussian FA, the MGP shrinkage prior (Bhattacharya and Dunson, 2011) defines an infinite sequence of loading columns: draw *δ*_1_ ~ Gamma(*a*_1_, 1), *δ*_*j*_ ~ Gamma(*a*_2_, 1) for *j* ≥ 2, set *γ*_*k*_ = Π_*j*≤*k*_ *δ*_*j*_, and place *W*_*d,k*_ | *γ*_*k*_, *α*_*d,k*_ ~ 𝒩 0, (*γ*_*k*_*α*_*d,k*_)^−1^ with *α*_*d,k*_ ~ Gamma(*ν/*2, *ν/*2). The construction admits an adaptive Gibbs sampler for inference that automatically learns the effective number of factors by pruning near-zero tail columns. Under mild conditions (e.g., 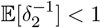, satisfied by *a*_2_ *>* 2 with unit rate), the truncation error decays exponentially with the truncation level, justifying finite truncations in practice.

### 3.2 CLING Model

We consider *M* data views 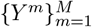 measured on the same *N* samples, where view *m* has *D*_*m*_ features. Let *K*_∞_ denote a (conceptually unbounded) number of latent factors shared across views via a common latent matrix 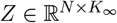, and view-specific loading matrices 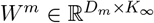. The likelihood is Gaussian with feature-wise heteroscedastic noise for *n* = 1, …, *N, m* = 1, …, *M, d* = 1, …, *D*_*m*_, *k* = 1, …, *K*_∞_:

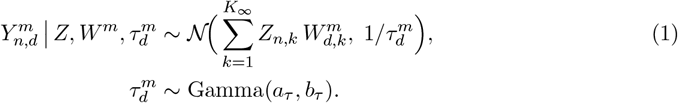

The latent factors are spherical:

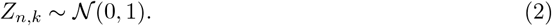

For each view *m*, we place a product-of-Gammas prior on the columns of *W*^*m*^, which induces progressively stronger column-wise shrinkage across *k* while remaining invariant to feature permutations; this promotes a finite effective number of factors and stable estimation in high-dimensional settings. Concretely, for *k* ≥ 1,

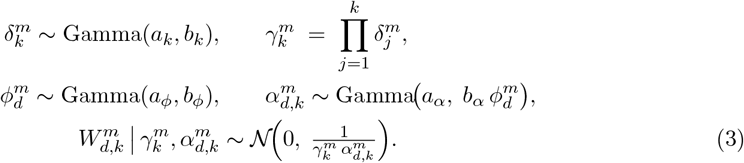

Here, at the view level, 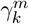 is a global column shrinker, whose expected log increases with *k* (cumulative shrinkage) whenever *ψ*(*a*_*k*_) *>* log *b*_*k*_ (where *ψ* denotes the digamma function), and 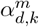 is a local feature–factor shrinker that controls the element-wise precisions of the loadings 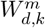, modulated by the row-specific 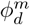.Together, this global–local hierarchical shrinkage on the loadings yields finite effective rank and preserves conjugacy for closed-form inference. The overall generative structure is summarized by the plate diagram in Figure 1.

**Figure 1.**
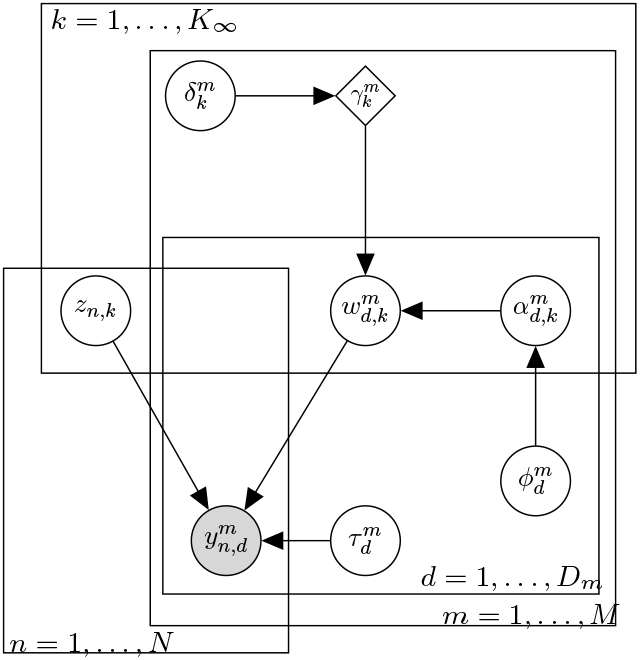
Graphical representation of CLING.

The cumulative shrinkage 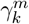 pushes the tail toward vanishing loadings, effectively forcing higher-index loading columns toward zero. Consequently, only a small subset of factors carries nontrivial posterior mass. Following Bhattacharya and Dunson (2011), the probability that the *K*_∞_-truncated covariance deviates from the infinite one by more than *ε >* 0 decays exponentially in *K*_∞_, so a modest *K*_∞_ suffices. Conceptually, the latent dimensionality is unbounded. In practice, we fix a large truncation level *K*_∞_, purely for computational convenience. This truncation does not preselect the number of factors: the global–local shrinkage automatically deactivates redundant columns, so the effective number of active factors is learned from the data rather than tuned.

### 3.3 Local prior specification

Element-wise precisions are governed by a Gamma–Gamma (*α*–*ϕ*) hierarchy, which preserves conjugacy while permitting flexible marginal specifications for *α* (concrete choices below).

#### Half-*t*/half-Cauchy cases

With *a*_*α*_ = *ν/*2, *b*_*α*_ = *ν*, and *a*_*ϕ*_ = 1*/*2, the local scale 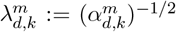 follows a 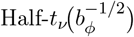 distribution (see Appendix A). For *ν* = 1, this specializes to half-Cauchy 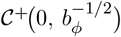; choosing *b*_φ_ = 1 yields the canonical 𝒞^+^(0, 1).

#### Beta–prime case

Marginalizing 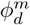 in the hierarchy yields a BetaPrime ( *a*_*α*_, *a*_*ϕ*_; scale 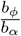) prior for the element-wise precision 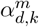 (see Appendix A).

#### Shrinkage structure

The variance of a loading is 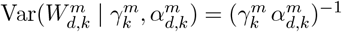, which acts as a soft gate on loadings. The local precision 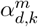 supplies feature–factor adaptivity within active columns: smaller 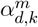 permits larger loadings, larger 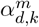 increases shrinkage. While the feature-level scale 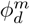 pools information across factors and steers the typical magnitude per feature (larger 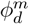 favors smaller 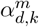 and greater signal retention; smaller 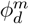 pushes mass toward larger 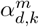 and stronger shrinkage). With heavy-tailed local-scale specifications arising from the Gamma–Gamma hierarchy, the prior exhibits selective shrinkage (an unbounded spike at zero with slab-like tails) robustly separating active vs. inactive entries. An illustrative visualization of the global and local shrinkage mechanisms acting on the loadings is provided in Appendix A.2 (Figures A.1–A.2).

### 3.4 Inference

Let ℋ:= {*a*_*τ*_, *b*_*τ*_, {*a*_*k*_, *b*_*k*_} _*k*≥1_, *a*_*ϕ*_, *b*_*ϕ*_, *a*_*α*_, *b*_*α*_} collect the hyperparameters. Conditional on ℋ, the joint density of CLING factorizes as

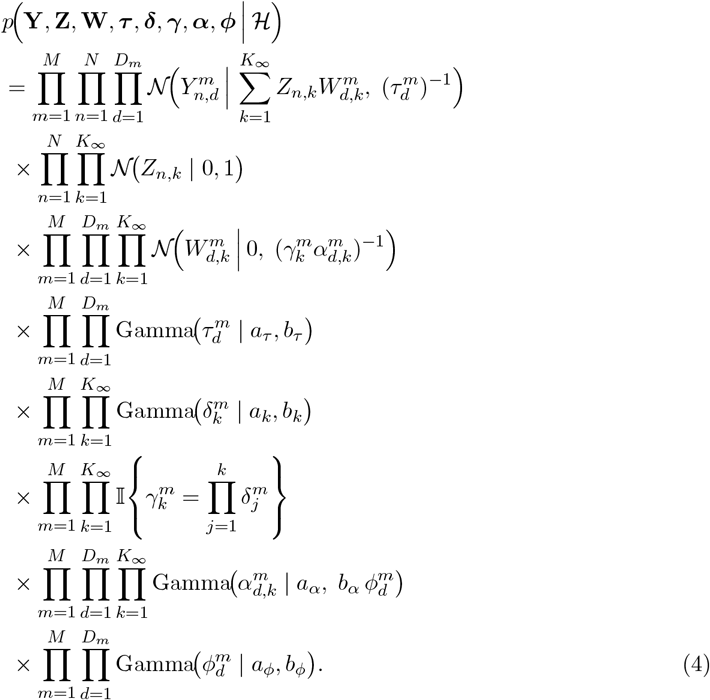

We fit the model with mean-field variational inference (MFVI) (Blei and Jordan, 2006; Doshi et al., 2009; Blei et al., 2017; Hansen et al., 2025) by maximizing the evidence lower bound (ELBO)

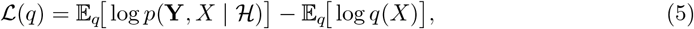

equivalently minimizing the KL divergence between the variational and exact posterior,

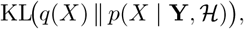

where *X* := {**Z, W, *τ***, ***δ, α, ϕ***} and 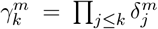 is deterministic. We use a structured mean-field factorization

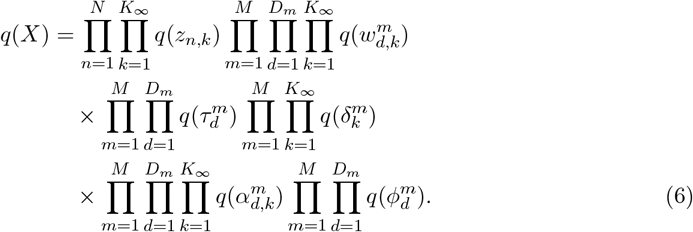

Under coordinate-ascent variational inference, the optimal variational distribution for any block *x*_*i*_ satisfies

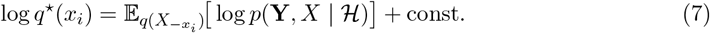

For the cumulative shrinkage updates, we have

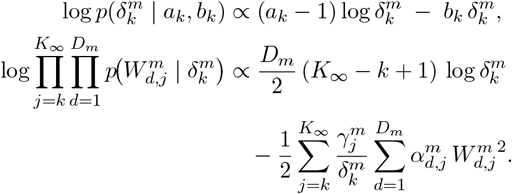

Applying (7) yields a Gamma variational posterior with parameters

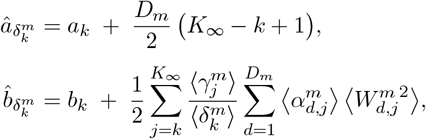

where 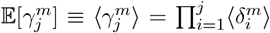. The remaining variational posteriors follow by conjugacy: Gaussian for *z*_*n,k*_ and 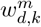, and Gamma for 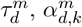, and 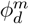 (all indices); see Appendix A for explicit expressions. We iterate until ELBO convergence.

#### Unsupervised factor number selection

To make CLING fully unsupervised, we adopt a prune/add schedule in the spirit of Bhattacharya and Dunson (2011). We initialize the truncation level as

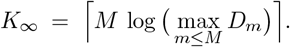

We then run variational inference and periodically assess the per-view variance explained by each factor. For view *m* and factor *k*, we compute

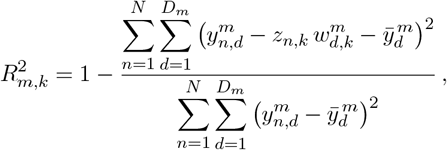

where 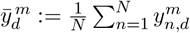 is the feature-wise mean used to center view *m*. Every 25 iterations (after a 100-iteration warm-up), we mark factor *k* inactive if 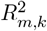 falls below a small threshold (typically 0.01–0.02) for all views *m*. If no factors are removed in a pruning step, we add a small batch of new factors. To discourage excessive oscillations, we allow at most three effective prune–after–add cycles; thereafter, we perform one final prune without adding new factors, freeze *K* at the resulting active count, and run an additional 50 iterations. This schedule complements the shrinkage prior and yields effective selection of the true number of factors without altering the variational objective.

## 4 Experiments

### 4.1 Simulations

To evaluate our model in a controlled setting where the ground truth is known, we first applied it on synthetic datasets and compared its performance against competing methods and ablation variants.

#### Simulation settings

We mimic the generative process typical to factor analysis with sparsity priors to create multi-view data using the following procedure: fix the number of latent factors *K* and draw the factor matrix *Z* ∈ ℝ^*N×K*^ with *Z*_*n,k*_ ~ 𝒩 (0, 1). For each view *m* = 1, …, *M*, feature *d* = 1, …, *D*_*m*_, and factor *k* = 1, …, *K*, sample a precision 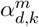 and first draw a presparsity loading 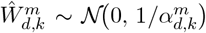. Impose elementwise sparsity with independent masks 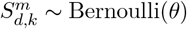, where *θ* = 1−sparsity, and set 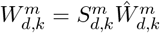. Finally, obtain observations by linearly combining factors with the resulting loadings and adding Gaussian noise with standard deviation *σ*, i.e., 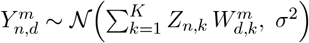.

As a baseline, we fixed the following values in our simulations: *N* = 100, *M* = 3, *D*_*m*_ = 400 for *m* = 1, 2, 3, *K* = 5, *σ*^2^ = 1, and sparsity level 1 − *θ* = 0.75. We then varied three parameters: the number of factors (*K* ∈ {1, 5, 10, 15, 20, 25}), the noise level (*σ* ∈ {0.5, 0.75, …, 2.0}), and the sparsity level ((1 − *θ*) ∈ {0.65, 0.70, …, 0.85}). The descriptions and results for different data generation settings are provided in Appendix B.

#### Evaluation

For each setting, we evaluated the methods using four criteria: (i) ability to infer the correct number of factors, (ii) the total explained variance (the higher, the better), (iii) the correspondence between true and inferred factors, measured as the average maximum Spearman correlation between each true factor and its closest inferred factor, and (iv) feature recovery, quantified as the average Jaccard index between the top 10% of true features (by absolute weight) and the features identified from the estimated weight matrices. Each experiment is repeated 20 times.

#### Competing methods and ablations

We compare our method against MOFA (Argelaguet et al., 2018), MuVI (without prior knowledge on weight matrices) (Qoku and Buettner, 2023), PCA (Pedregosa et al., 2011), and Tucker decomposition (Kossaifi et al., 2019). Among these, only MOFA explicitly removes inactive factors. For the other methods, we apply a post-training threshold of 0.01 variance explained per factor to distinguish active from inactive factors. For multiview data, Tucker decomposition requires the input to be an *N* × *M* × *D* tensor, which means all views must have the same number of features. In addition, we evaluate two ablation variants: (i) CLING_MGP_, which ablates the Gamma-Gamma hierarchy and retains a single local Gamma(1.5, 1.5) prior, i.e., a straightforward multi-view extension of the MGP. (ii) CLING_ARD_: cumulative shrinkage is applied only across factors; the local precision is removed and replaced by a view-level precision with a Gamma(1.5, 1.5) prior, analogous to ARD in MOFA. All remaining hyperparameters are set to the same values as in CLING (see Appendix B).

#### Results

Across a wide range of sparsity and noise levels (Figure 2, top row), CLING accurately recovers the true number of latent factors. Most competing methods fail under high noise or high sparsity; among them, MOFA is comparatively more robust in these settings but frequently breaks down as *K* increases, exhibiting instability and tending to under/over-estimate *K*. Furthermore, CLING outperforms all compared methods, in recovering the true latent factors and the sets of most important features for each factor (Figure 2, 3rd and 4th row), yielding the most identifiable solution without requiring any additional prior knowledge. At the same time, comparable or higher total explained variance (Figure 2, 2nd row) demonstrates that CLING captures at least as much variability as the other methods. These results remain consistent across diverse simulation settings, including scenarios that do not conform to CLING’s modeling assumptions (see Appendix, Figure B.3 and B.4). We note that although both CLING and MOFA rely on an explained variance threshold, CLING is more stable in the number of factors selected across different threshold values (Appendix, Figure B.5).

**Figure 2.**
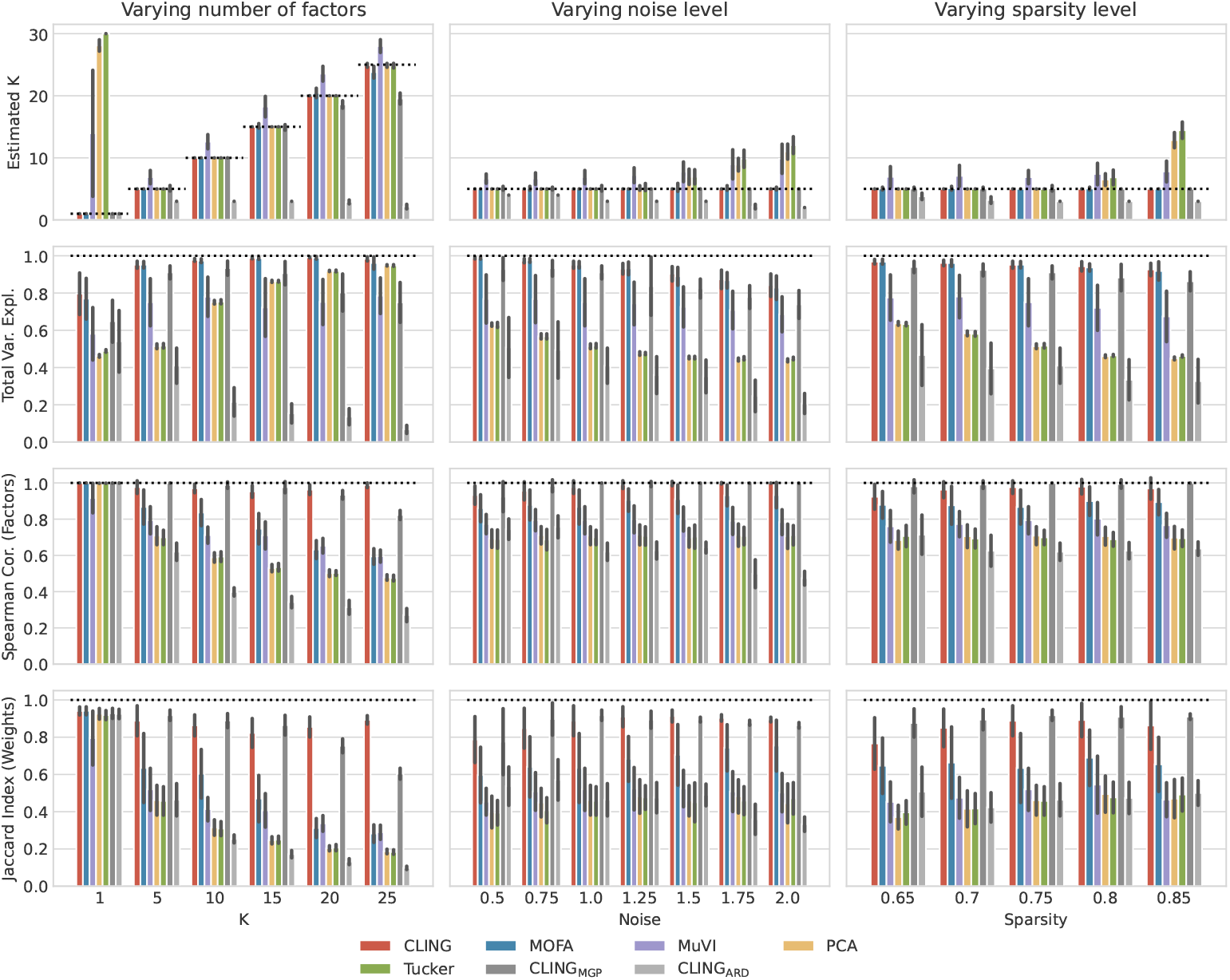
Comparison of true versus inferred number of factors, latent factor, and weights, and total explained variance for CLING, four competing models, and three ablation models across varying numbers of factors, noise levels, and sparsity levels. The bars show mean ± standard deviation. Dashed lines indicate optimal performance.

**Figure 3.**
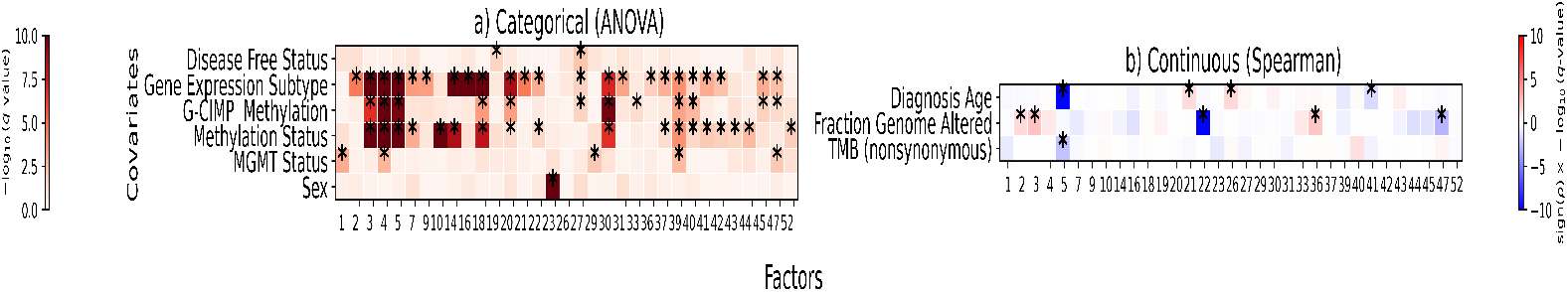
Associations between inferred latent factors (x-axis) and sample covariates (y-axis). For categorical covariates (a), colors indicate − log_10_(*q* − value); for continuous covariates (b), colors indicate sign(*ρ*) × −log_10_(*q* − value), where *ρ* is the Spearman correlation. Asterisks mark significant associations (*q* = 0.05).

Compared with the ablations, the full CLING model achieves superior performance across all evaluation metrics. This advantage likely stems from the Gamma-Gamma hierarchy, which counteracts the tendency of cumulative shrinkage to become excessively strong as the number of active factors grows. In the single-Gamma ablation (CLING_MGP_), the model systematically underestimates *K* once *K >* 15 and performs markedly worse, whereas CLING preserves the true K. By contrast, replacing the local precision with a view-level precision and applying cumulative shrinkage only across factors CLING_ARD_ yields overly aggressive shrinkage and uniformly poor results.

### 4.2 Predictive Performance on Biological Datasets

Next, we evaluated the ability of CLING to infer meaningful latent factors from heterogeneous multiview data and to support downstream predictive tasks.

#### Datasets

We analyzed three biological datasets: (i) Developmental bulk RNA-seq across species and organs (**Evo-Devo** dataset), spanning multiple organs for different species and a range of developmental age values, obtained from the *MEFISTO* study (Velten et al., 2022). For this dataset, five separate views were defined by organs. Predicted covariates included developmental age and species. (ii) Early gastrulation single-cell multi-omics data measured using single-cell nucleosome, methylation and transcription sequencing (**scNMT-seq** data) (Argelaguet et al., 2019) in mouse embryos at different stages of gastrulation for various lineages. For scNMT-seq data, three views were defined by modalities. Covariates included cell type and developmnental stage. (iii) Glioblastoma multiforme (GBM) tumor samples from the The Cancer Genome Atlas (Gao et al., 2013) (**GBM** dataset; Brennan et al. (2013)), with two bulk sequencing views: gene expression and DNA methylation. Covariates included tumor gene expression subtype, G-CIMP and global methylation status, MGMT promoter status, disease-free status, patient sex, age at diagnosis, fraction genome altered, and tumor mutational burden (TMB). Details of predicted covariates and normalization for each dataset are in Appendix C.

#### Evaluation metrics

To quantify the quality of the inferred factors for each dataset, we trained random forest regressors and classifiers to predict sample-level covariates from the factors in a 5-fold cross-validation scheme. To assess predictive performance, we used *R*^2^ for regression and F1 score for classification tasks.

#### Compared methods

CLING was compared to MOFA (Argelaguet et al., 2018) (as the top performing multi-view model based on simulations) and PCA (Hotelling, 1933) as a simple baseline that can handle non-identical feature sets across views.

#### Results

As shown in Table 1, for the **Evo-Devo** dataset, CLING achieved the highest *R*^2^ for developmental age prediction while all methods achieved perfect species classification, demonstrating that CLING recovers temporally coherent and cross-species factors. For the **scNMT-seq** data, as presented in Table 2, CLING outperformed compared methods in three out of four tasks defined by prediction of developmental stage and lineage-specific markers. These results demonstrate that CLING effectively integrates heterogeneous modalities, capturing both shared developmental signals and modality-specific variation. Finally, for the **GBM** dataset CLING factors achieved superior predictive performance in predictive disease-related covariates, such as clinical subtype and methylation status, suggesting that they encode biologically relevant information associated with GBM heterogeneity (Table 3).

**Table 1.** Predictive performance (mean scores ± standard deviations) for the Evo-Devo dataset.

| Covariate | CLING | MOFA | PCA |
| --- | --- | --- | --- |
| Age ($R^2$) | **0.949 ± 0.047** | 0.921 ± 0.041 | 0.939 ± 0.053 |
| Species (F1) | 1.000 ± 0.000 | 1.000 ± 0.000 | 1.000 ± 0.000 |

**Table 2.**
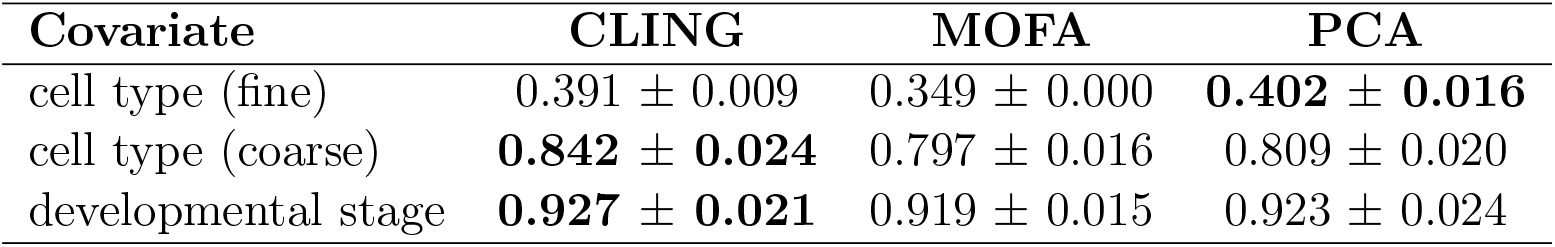
Predictive performance (mean F1 scores ± standard deviations) for the scNMT-seq dataset.

| Covariate | CLING | MOFA | PCA |
| --- | --- | --- | --- |
| cell type (fine) | 0.391 ± 0.009 | 0.349 ± 0.000 | **0.402 ± 0.016** |
| cell type (coarse) | **0.842 ± 0.024** | 0.797 ± 0.016 | 0.809 ± 0.020 |
| developmental stage | **0.927 ± 0.021** | 0.919 ± 0.015 | 0.923 ± 0.024 |

**Table 3.** Predictive performance (mean F1 scores ± standard deviations) for the GBM dataset.

| Covariate | CLING | MOFA | PCA |
| --- | --- | --- | --- |
| Disease Free Status | **0.514 ± 0.050** | 0.467 ± 0.044 | 0.445 ± 0.047 |
| G-CIMP Methylation | **0.916 ± 0.110** | 0.902 ± 0.118 | 0.815 ± 0.079 |
| Gene Expression Subtype | **0.780 ± 0.044** | 0.778 ± 0.066 | 0.762 ± 0.042 |
| MGMT Status | 0.546 ± 0.041 | **0.614 ± 0.039** | 0.609 ± 0.073 |
| Methylation Status | **0.781 ± 0.074** | 0.764 ± 0.077 | 0.615 ± 0.047 |
| Sex | **0.991 ± 0.013** | 0.988 ± 0.012 | 0.941 ± 0.017 |

### 4.3 CLING Interpretability on brain cancer data

Finally, to showcase the ability of CLING to interpret the analyzed data and generate testable biological hypotheses, we assessed individual factor associations with sample covariates and identified pathways enriched in respective loadings for the GBM dataset. For all statistical tests, resulting *p*-values were adjusted for multiple testing using the Benjamini–Hochberg correction, with significance threshold of *q* = 0.05.

#### Factor-covariate associations

CLING inferred 54 latent factors for GBM data. To assess the association of each factor with continuous sample covariates, we computed Spearman correlations. For categorical covariates, associations were tested using one-way ANOVA.

As shown in Figure 3a, the covariate that the most factors are associated with is Gene Expression Subtype (with values Proneural, Neural, Classical, and Mesenchymal, and G-CIMP, where the former four are related to transcriptional, while the latter to methylation patterns of tumors). These subtypes differ in their genomic features and response to therapies (Verhaak et al., 2010). Thus, they are expected to be the main source of variation in GBM data, confirming the biological relevance of CLING’s factors. The second most commonly associated covariate is Methylation Status, highlighting the importance of this genomic alteration for GBM. F5 is among factors associated with covariates Gene Expression Subtype, G-CIMP Methylation, Methylation and MGMT Status, indicating that it captures shared mechanisms between these clinical features. Figure 4 displays monotonic decrease of F5 from methylation status C1 to C4, with higher values obtained for C6 and G-CIMP. At the same time, F5 is decreased in the Classical and increased in G-CIMP subtypes. Apart from F5, other factors of interest were F4 (also associated with multiple categories), F10 (exclusively strongly associated with Methylation Status), F16 (exclusively associated with Gene Expression Subtype), F23 (associated with sex) and F27 (most strongly associated with Disease Free Status). For numerical covariates (Figure 3b), strong associations were found again for factor F5 with Diagnosis Age and for Fraction Genome Altered (F22).

**Figure 4.**
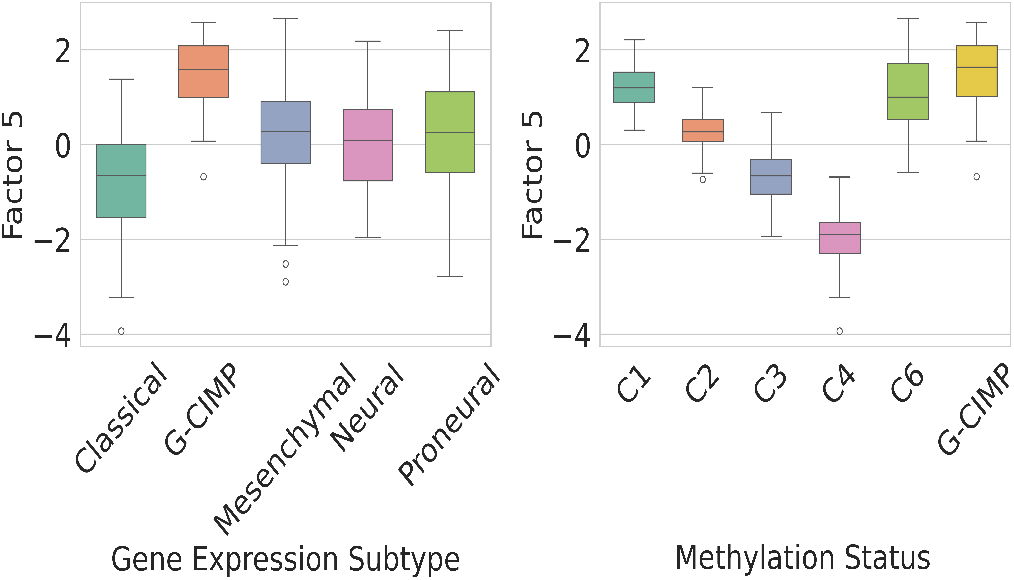
Distribution of CLING latent factor F5 values across covariate classes gene expression subtypes and methylation status of samples. Each boxplot corresponds to a factor–covariate category pair.

##### Factor-pathway enrichment

To further interpret the factors of interest (F4, F5, F10, F16, F22, F23, F27), we performed gene set enrichment analysis (GSEA; Subramanian et al. (2005)) on the gene expression loadings of these factors. For each factor, genes (or features for each view) were ranked by their factor weights, and enrichment was assessed against Hallmark 50 gene sets representing core biological pathways (Liberzon et al., 2019). Enrichment scores were computed using the running-sum statistic, and empirical *p*-values were estimated via 10,000 permutations of gene labels to establish a null distribution.

This analysis revealed the relevance of known gene set pathways on GBM covariates associated to the factors (Figure 5), demonstrating that CLING captures latent biological processes that are both interpretable and predictive. Recall that factor F5 is positively associated with C1 and C6 statuses, while showing inverse associations with the Classical subtype and C4 status, as well as a negative correlation with patient age. Functionally, gene expression loadings for F5 are negatively enriched for DNA repair, G2-M checkpoint, and MYC target pathways, indicating increase of proliferative and repair-related programs in older patients and the Classical subtype. In contrast, factor F4, with lower values in G-CIMP tumors (Appendix, Figure C.6), exhibits enrichment in the opposite direction to F5 for E2F targets and G2-M checkpoint pathways, highlighting transcriptional programs specific to G-CIMP tumors.

**Figure 5.**
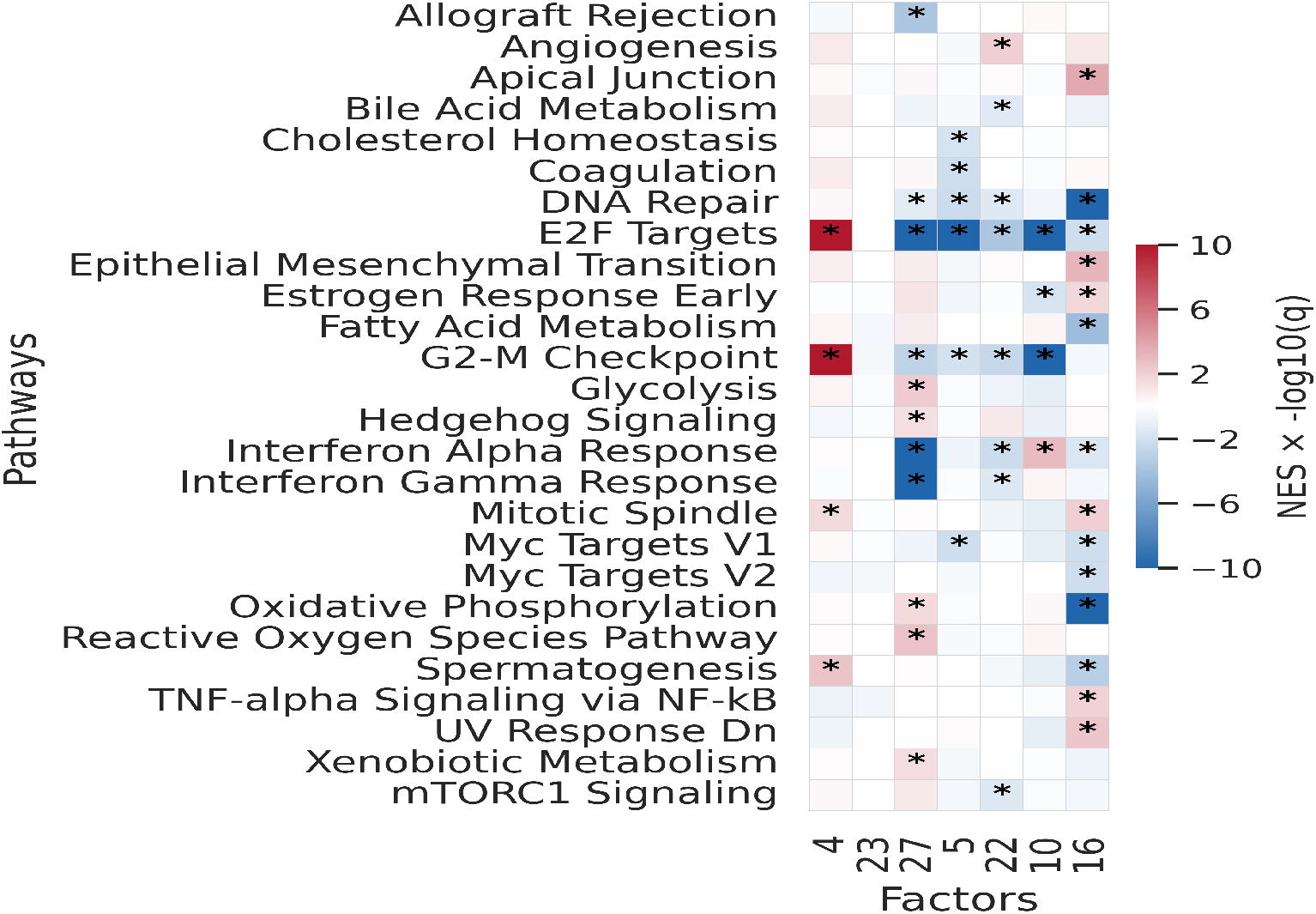
GSEA of factor loadings across biological pathways. Colors indicate positive (red) or negative (blue) association of pathways (y-axis) with factor loadings (x-axis). Stars indicate statistically significant associations.

## 5 Conclusions

We introduced CLING, an unsupervised, conjugate multi-view factor model that combines cumulative column-wise shrinkage with a Gamma-Gamma local-precision hierarchy to enable automatic factor selection and precise estimation of loadings. Across synthetic and biological datasets, CLING recovered more accurate factors and informative loadings, with comparable or higher explained variance than strong baselines; ablations confirmed that the Gamma-Gamma hierarchy stabilizes cumulative shrinkage as *K* grows. Results on real data established that CLING effectively integrates multiview data, uncovers latent structure reflecting underlying biology, and produces factors that are useful for both predictive modeling and mechanistic interpretation across diverse datasets.

Looking ahead, further gains may come by extending CLING to non-Gaussian likelihoods, integrating topic-model components to handle structured views alongside simple views, and introducing further nonparametric elements, such as an Indian Buffet Process prior for factor usage and mixture components for clustering. Together with closed-form variational updates, these directions position CLING as a robust default for large, heterogeneous multi-omics integration.

## Supplementary Information

### A Foundations and Inference for CLING

This appendix complements the main text by outlining the marginals under the Gamma–Gamma hierarchy (half-*t*/half-Cauchy on local scales *λ* = *α*^−1*/*2^; Beta–prime on precisions *α*), specifying the mean-field variational family, deriving closed-form coordinate updates, and presenting the ELBO used for convergence monitoring, with additional illustrative examples provided for intuition.

#### A.1 Half-*t* scales; Beta-prime precisions

Below we show that the marginal prior of 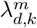 is Half-*t*_*ν*_ with scale 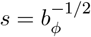; in particular, for *ν* = 1 it specializes to a half-Cauchy with the same scale.

##### Proposition 1

Let | *α ϕ* ~ Gamma(*ν/*2, *ν ϕ*), *ϕ* ~ Gamma(1*/*2, *b*_*ϕ*_), and set *λ* = *α*^−1*/*2^. Then the marginal density of *λ* is

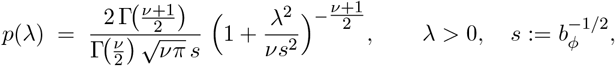

i.e., *λ* ~ Half-*t*_*ν*_ (*s*). For *ν* = 1, this reduces to the half-Cauchy density

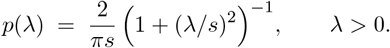

**Proof**. Conditioning on *ϕ*, the density of *α* is

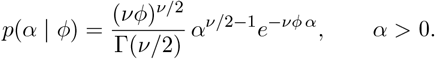

Apply the change of variables *α* = *λ*^−2^ with Jacobian d*α/*d*λ* = 2 *λ*^−3^ to obtain

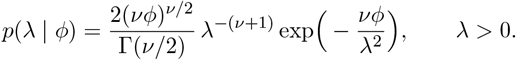

Marginalize *ϕ*:

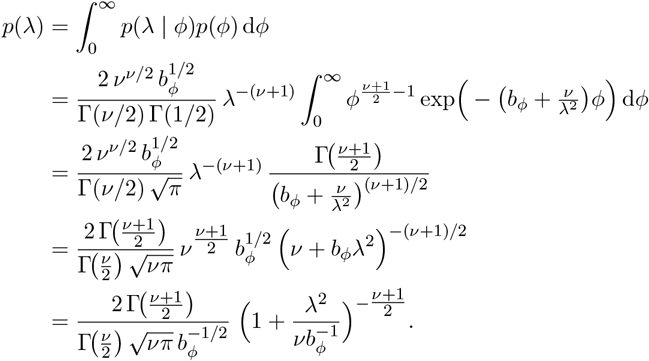

Recognizing 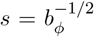 yields the stated Half-*t*_*ν*_ (*s*) density. For *ν* = 1, this specializes to the half-Cauchy 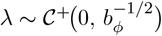; in particular *b*_*ϕ*_ = 1 gives the canonical 𝒞^+^(0, 1).

##### Proposition 2

Let *α* | *ϕ* ~ Gamma(*a*_*α*_, *b*_*α*_ *ϕ*) and *ϕ* ~ Gamma(*a*_*ϕ*_, *b*_*ϕ*_). Then the marginal distribution of element-wise precision *α* is BetaPrime *a*_*α*_, *a*_*ϕ*_; scale 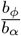, with density for *x >* 0,

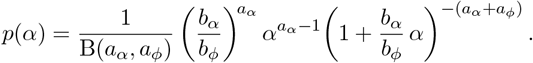

**Proof**. Marginalize *ϕ*:

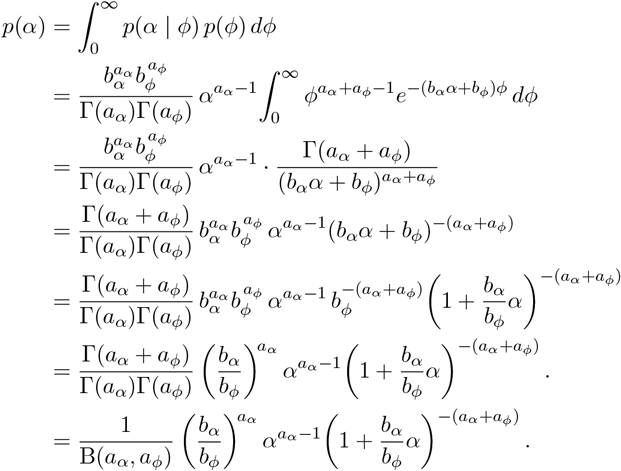

This is the Beta–prime density with scale *b*_*ϕ*_*/b*_*α*_.

#### A.2 Illustration of the generative shrinkage mechanism

We first provide an illustrative example of the shrinkage mechanisms acting on the loading matrices in CLING, using two views, to build intuition for how hierarchical global and local shrinkage induces sparsity.

**Figure A.6.**
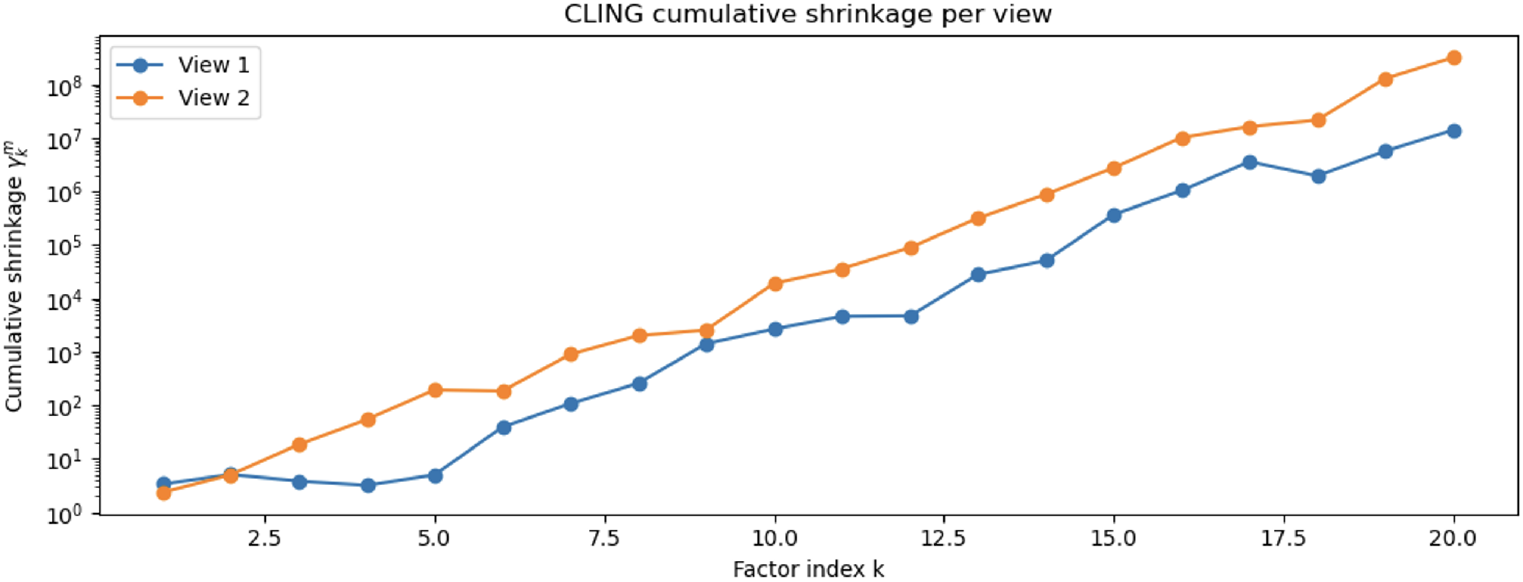
Global shrinkage values in CLING. The plotted cumulative shrinkage values 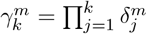, which increase with *k* and progressively shrink higher indexed factors toward zero.

**Figure A.7.**
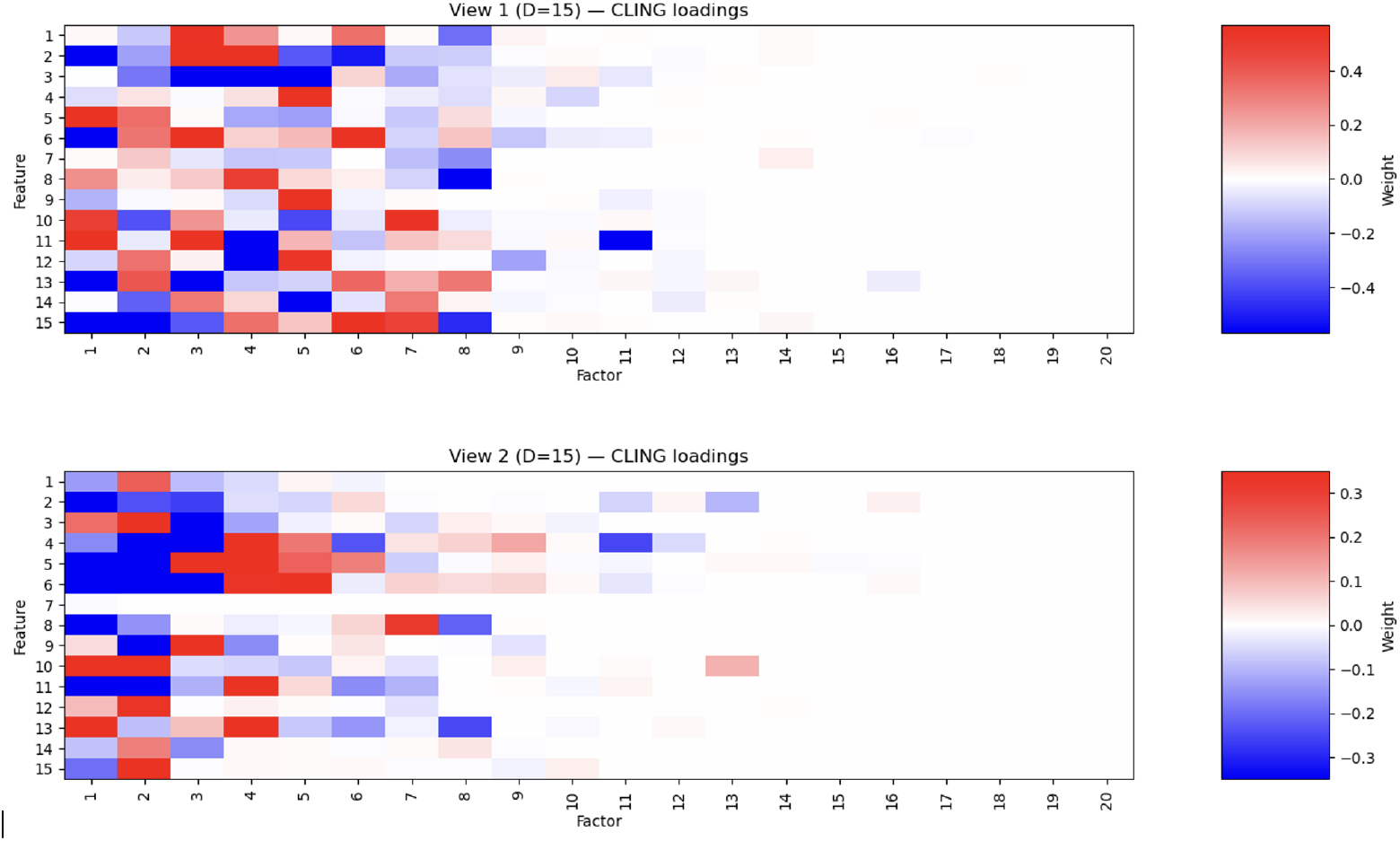
Global-local shrinkage of loadings (two views). Each loading 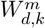 is controlled by a global factor shrinkage 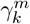 and a local precision 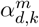, inducing view-specific sparsity patterns.

#### A.3 Coordinate-Ascent Variational Inference (CAVI) Updates

All updates are mean-field CAVI steps.

##### Scores *Z*

For each (*n, k*),

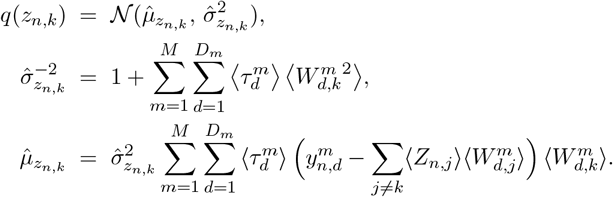

##### Loadings *W*

For each (*m, d, k*),

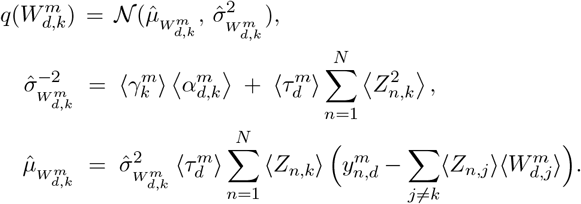

##### Noise precisions *τ*

For each (*m, d*),

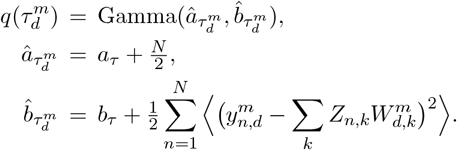

##### Local precisions *α*

For each (*m, d, k*),

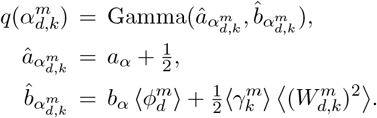

##### Feature scales *ϕ*

For each (*m, d*),

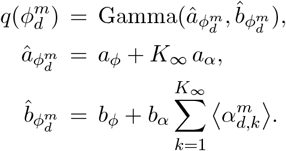

#### A.4 Evidence Lower bound

We monitor convergence by the variational evidence lower bound (ELBO), which we decompose into a likelihood term and one term per latent variable.

##### Mean-field factorization

With *X* := {**Z, W, *τ***, ***δ, α, ϕ***}, the ELBO decomposes as

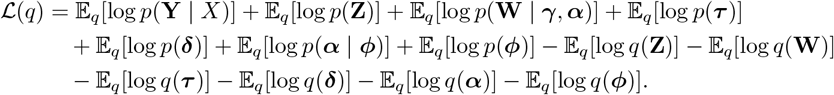

- **Gaussian likelihood term**:

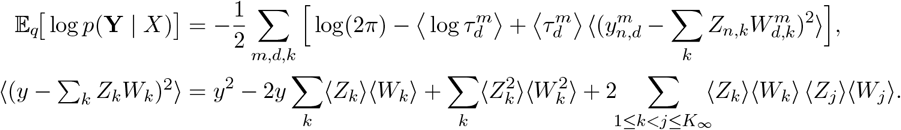
- **Z prior and entropy terms**:

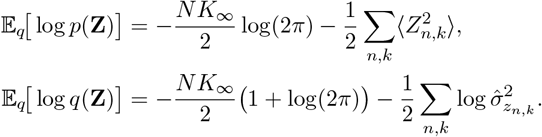
- **W prior and entropy terms**:

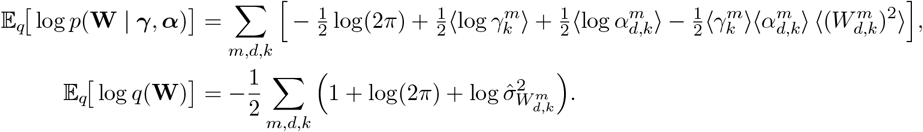
- ***τ* prior and entropy terms**:

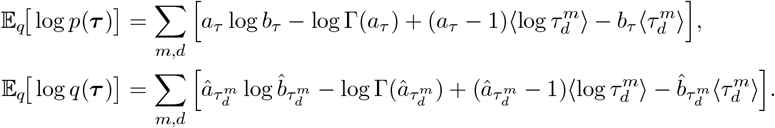
- ***δ* prior and entropy terms**:

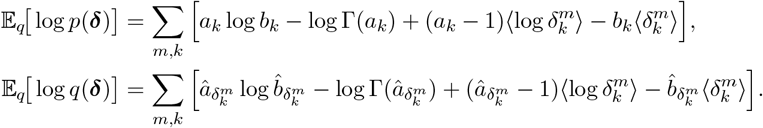
- ***α* prior and entropy terms**:

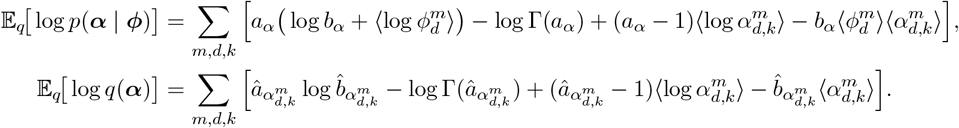
- ***ϕ* prior and entropy terms**:

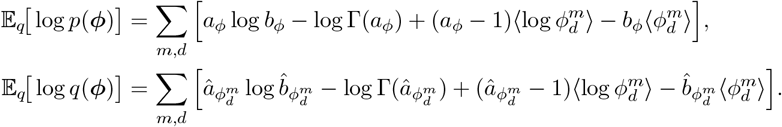

### B Simulations

#### B.1 Simulation overview

In our simulator, described in the main text, we intentionally depart from CLING’s model assumptions by adopting a MOFA-style sparsity mechanism: we suppress cumulative column shrinkage by fixing 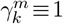 for all *m, k* and induce sparsity via an element-wise Bernoulli mask on the loadings. In addition, we replace MOFA’s column-level ARD 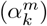 with fully local ARD precisions 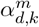 (per view–feature–factor). In this way, we induce stronger and more heterogeneous sparsity patterns than introduced by MOFA simulator.

#### B.2 Supplementary simulations

We conduct three supplementary simulations. Figure B.1 targets a high-dimensional, higher-noise regime (baseline true number of factors K=10); Figure B.2 repeats the analysis in a moderate-dimensional, lower-noise setting (baseline K=5); and Figure B.3 examines sensitivity to the pruning threshold for factor activity, comparing CLING and MOFA across several thresholds.

#### B.3 Model configuration

##### CLING configuration

Unless stated otherwise, we use *a*_*τ*_ =*b*_*τ*_ =10^−3^; cumulative-shrinkage Gammas *a*_1_=2.0, *b*_1_=1.0, *a*_*k*_=2.1, *b*_*k*_=1.0 for *k* ≥ 2; Gamma–Gamma local precisions with *a*_*α*_=1*/*2, *b*_*α*_=1, *a*_*ϕ*_=1*/*2, and *b*_*ϕ*_=1(half-Cauchy local scales).

##### MOFA configuration

We used `mofapy2` v0.7.2 (https://github.com/bioFAM/mofapy2). Views were centered to match CLING’s preprocessing. We enabled weight-level sparsity via `ard_weights=True` and `spikeslab_weights=True`, and disabled factor-level sparsity (`ard_factors=False`, `spikeslab_factors=False`). The maximum number of factors was set to *K*=30 in all simulations. During training, factor pruning used MOFA’s 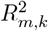 drop threshold `dropR2=0.01` by default; for Fig. B.3 we varied the threshold across {0.02, 0.01, 0.005}.

##### MuVI configuration

We used `MuVI` v0.1.5 (https://github.com/MLO-lab/MuVI) with `prior_masks=None` and maximum factors *K*=30. Data were normalized using MuVI’s default (centering and scaling by the global standard deviation). After training, we pruned factors whose per-factor explained variance fell below 0.01.

##### PCA configuration

We ran PCA with *K*=30 components using `scikit-learn` v1.7.1. Data were standardized in the usual way (feature-wise centering and scaling to unit variance). After training, components with per-factor explained variance fraction *<* 0.01 were discarded.

##### Tucker configuration

We applied a Tucker model with multilinear rank [*D, M, K*] and *K*=30 using `tensorly` v0.9.0. Data were standardized in the same manner as PCA. After training, we retained only components whose per-factor explained variance exceeded 0.01.

**Figure B.8.**
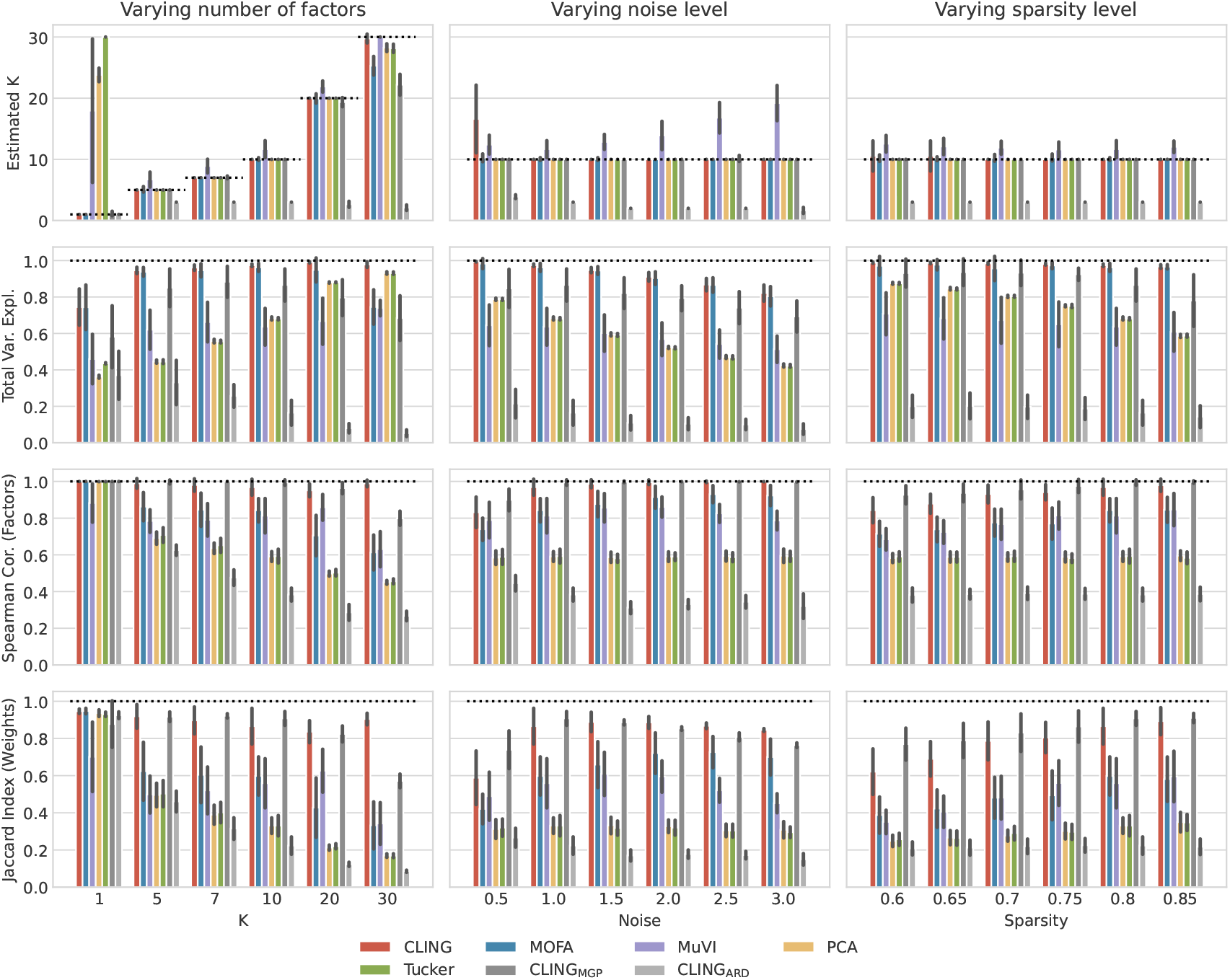
High-dimensional, high-noise scenario (baseline *K* = 10). Comparison of true versus inferred number of factors, latent factor, and weights, and total explained variance for CLING, four competing models, and two ablation models across varying numbers of factors, noise levels, and sparsity levels. The bars show mean ± standard deviation. Dashed lines indicate optimal performance. **Simulation settings** Data was generated as described in the main text. The baseline parameters are: *N* = 100, *M* = 3, *D*_*m*_ = 1000 for *m* = 1, 2, 3, *K* = 10, *σ* = 1, and sparsity level 0.8.

**Figure B.9.**
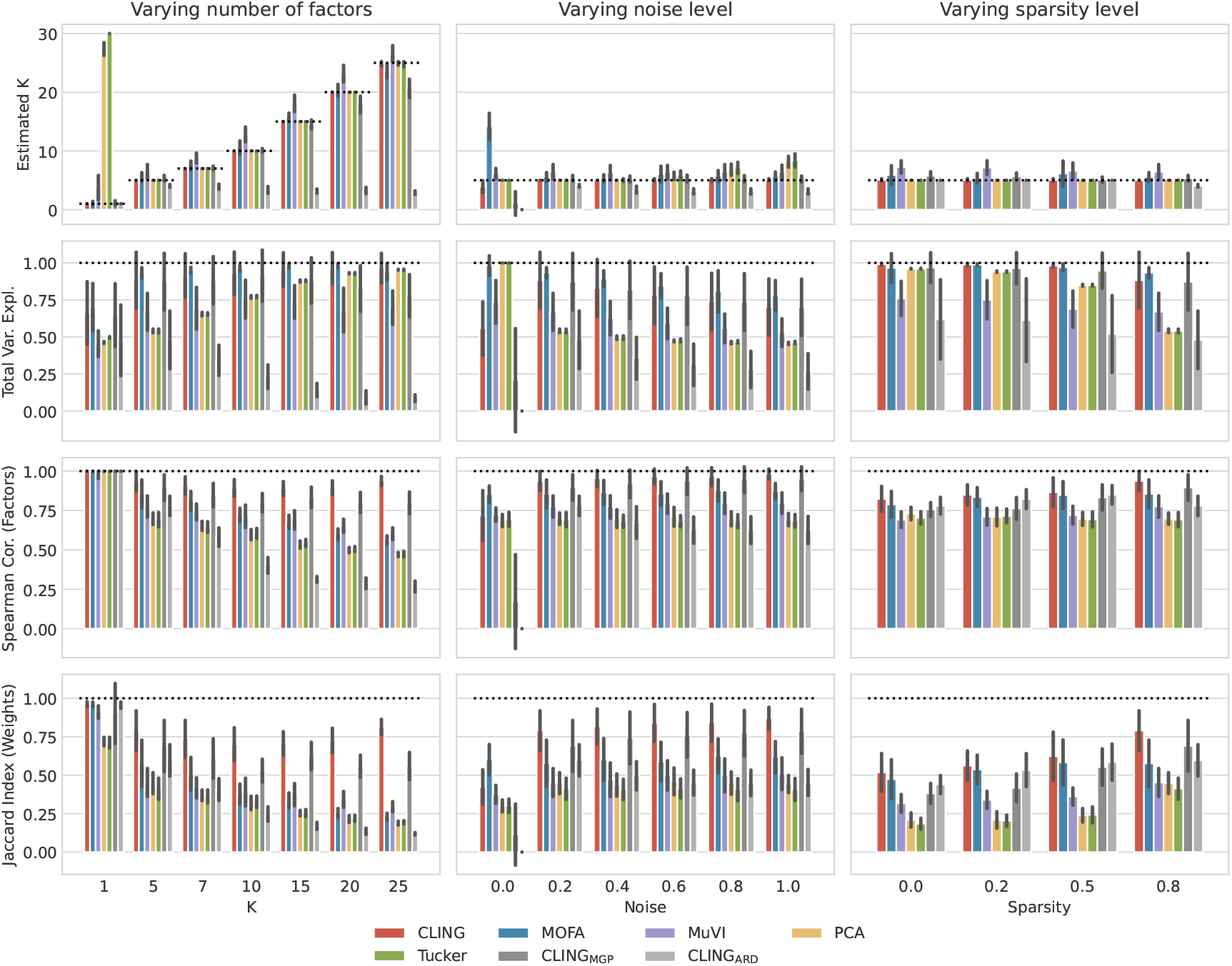
Moderate-dimensional, low-noise scenario (baseline *K* = 5). Comparison of true versus inferred number of factors, latent factor, and weights, and total explained variance for CLING, four competing models, and two ablation models across varying numbers of factors, noise levels, and sparsity levels. The bars show mean ± standard deviation. Dashed lines indicate optimal performance. **Simulation settings** Data was generated as described in the main text. The baseline parameters are: *N* = 100, *M* = 3, *D*_*m*_ = 400 for *m* = 1, 2, 3, *K* = 5, *σ* = 0.2, and sparsity level 0.8.

**Figure B.10.**
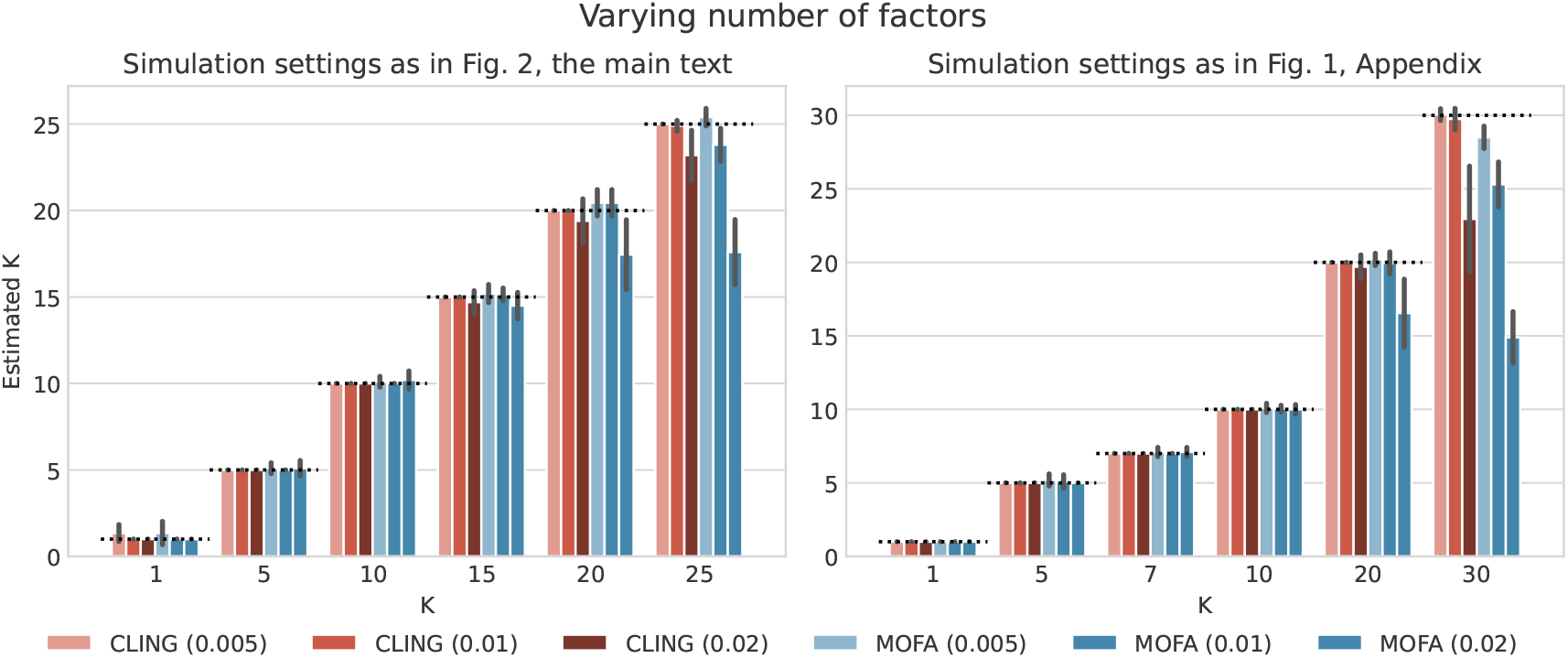
Pruning-threshold sensitivity (CLING vs. MOFA). Sensitivity of CLING and MOFA to pruning thresholds on explained variance (tresholds in {0.005, 0.01, 0.02}) in inferring the number of factors. The bars show mean ± standard deviation. Dashed lines indicate optimal performance. **Simulation settings** We used the same simulation settings as in other figures: Left panel - as in Fig. 2 of the main text; Right panel - as in Fig. B.1 in the Appendix.

### C Biological data analyses

#### C.1 Data preprocessing

All datasets were processed through a standardized multimodal integration pipeline designed to ensure comparability across biological contexts. Each dataset consists of one or more views, corresponding to distinct molecular modalities. The preprocessing and model initialization steps followed a unified protocol:

- **Missing values:** Missing entries in molecular views were replaced by zeros. Although CLING and MOFA can handle missing data natively, methods such as PCA, MuVI, and Tucker require complete input matrices.
- **Normalization:** Each view was independently *z*-score normalized to center and scale features.
- **Feature selection:** To reduce dimensionality and emphasize signal-bearing features, only features with high variance were used.

#### C.2 Evo-Devo dataset

The **Evo-Devo dataset** comprises transcriptomic measurements across five organs (brain, cerebellum, heart, liver, and testis) and five species (human, opossum, mouse, rat, and rabbit) sampled at multiple developmental time points Cardoso-Moreira et al. (2019). Each organ represents a distinct molecular view (*V* = 5). After filtering to one-to-one orthologs across species, each view contained 7,696 genes, with 500 highly variable genes retained per view.

**Table C.4.** Summary of species-specific developmental covariates in the Evo-Devo dataset. “Time” represents the developmental stage index (1–23) spanning embryonic to adult stages. For each species, five organs (Brain, Cerebellum, Heart, Liver, and Testis) contribute one view each.

| Species | N Samples | Developmental time (range) |
| --- | --- | --- |
| Human | 23 | 1 - 23 |
| Mouse | 14 | 1 - 14 |
| Opossum | 15 | 1 - 15 |
| Rabbit | 15 | 1 - 15 |
| Rat | 16 | 1 - 16 |

#### C.3 scNMT dataset

The **scNMT dataset** comprises single-cell multi-omics measurements collected during mouse gastrulation **?**, with 1,518 cells represented across three molecular views: RNA expression, DNA methylation, and chromatin accessibility (500 highly variable features per view).

#### C.4 GBM dataset

The **Glioblastoma Multiforme (GBM) dataset** integrates gene expression and DNA methylation data from 360 patients Brennan et al. (2013). Two molecular views were considered: RNA expression and DNA methylation (5,000 features each).

**Table C.5.**
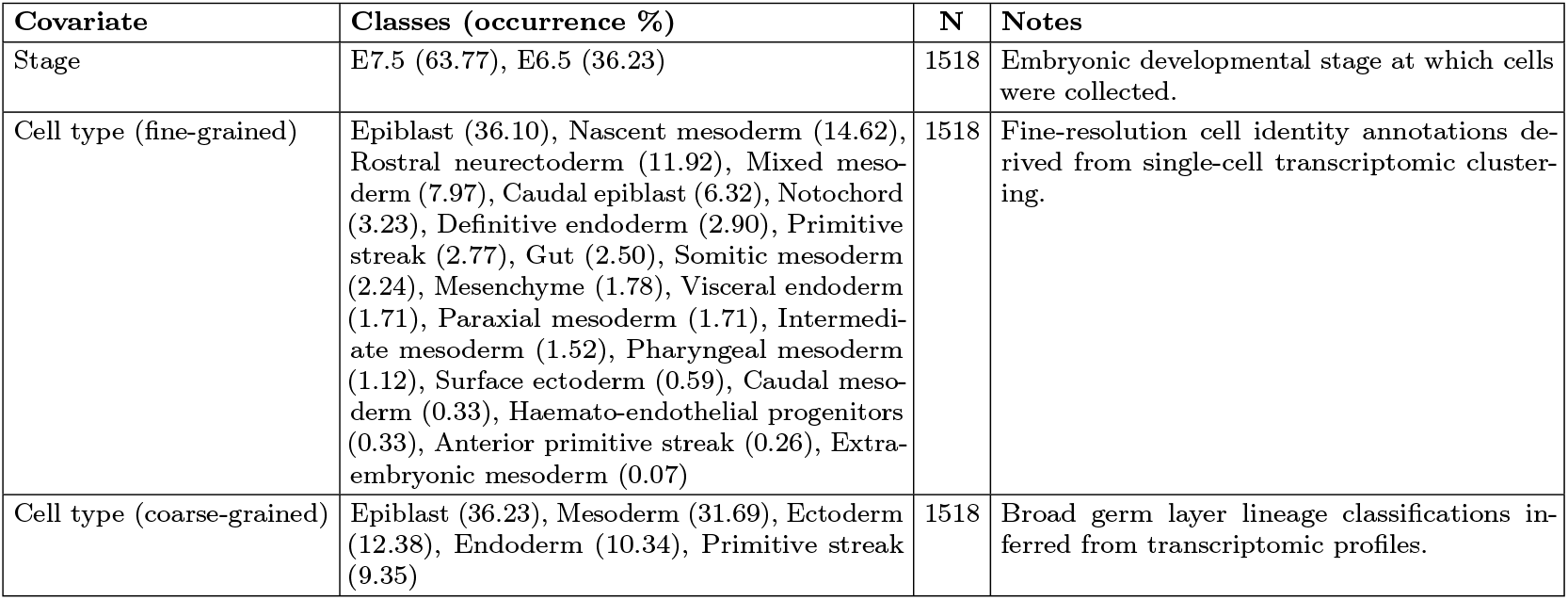
Classification-type covariates and class proportions for the mouse gastrulation dataset. “N” denotes the number of samples with non-missing data. “Classes” lists the categorical levels within each covariate and their relative frequencies (%).

**Table C.6.** Regression-type covariates and summary statistics for the GBM dataset. “N” indicates samples with non-missing data. “Summary Statistics” indicates the range, mean and standard deviation of the covariate values.

| Covariate | Summary Statistics | N | Notes |
| --- | --- | --- | --- |
| Diagnosis Age | Range = [10.9, 89.3]; Mean = 58.93; SD = 14.38 | 353 | Age at initial pathologic diagnosis. |
| Fraction Genome Altered | Range = [0.000, 0.795]; Mean = 0.213; SD = 0.131 | 343 | Fraction of the genome showing copy number alterations. |
| TMB (nonsynonymous) | Range = [0.100, 3.833]; Mean = 1.393; SD = 0.491 | 206 | Tumor mutational burden (nonsynonymous mutations per megabase). |

**Table C.7.** Classification-type covariates and category proportions for the GBM dataset. “N” indicates samples with non-missing data. “Classes” indicates the classes inside each covariate and their percentage of occurrences in non-missing data.

| Covariate | Classes (occurrence %) | N | Notes |
| --- | --- | --- | --- |
| Disease Free Status | Recurred/Progressed (60.17), DiseaseFree (39.83) | 354 | Indicates whether the patient remained disease-free at last follow-up. |
| Gene Expression Subtype | Mesenchymal (30.23), Classical (27.12), Proneural (18.64), Neural (16.10), G-CIMP (7.91) | 354 | Consensus GBM transcriptional subtypes. |
| G-CIMP Methylation | non-G-CIMP (91.53), G-CIMP (8.47) | 354 | Glioma CpG Island Methylator Phenotype; distinguishes IDH-mutant from wild-type tumors. |
| Methylation Status | C3 (27.59), C2 (24.71), CL4 (14.66), C1 (13.79), C6 (10.92), G-CIMP (8.33) | 348 | Global or locus-specific methylation classification. |
| MGMT Status | Unmethylated (51.01), Methylated (48.99) | 347 | MGMT promoter methylation; predicts response to alkylating chemotherapy. |
| Sex | Male (58.64), Female (41.36) | 353 | Biological sex of the patient. |

#### C.5 Distribution of latent factors across clinical covariates

#### C.6 Model configuration

##### CLING configuration

The CLING model was initialized with the number of latent factors set to *M* log(max_*m*≤*M*_ *D*_*m*_), where *M* is the number of views and *D*_*m*_ is the number of features in view *m*. The model was trained for up to 1000 iterations with an explained-variance threshold of 0.01, using the prune/add schedule described in the main text; the resulting active factor count *K*_final_ reflects the model’s adaptive complexity for each dataset.

**Figure C.11.**
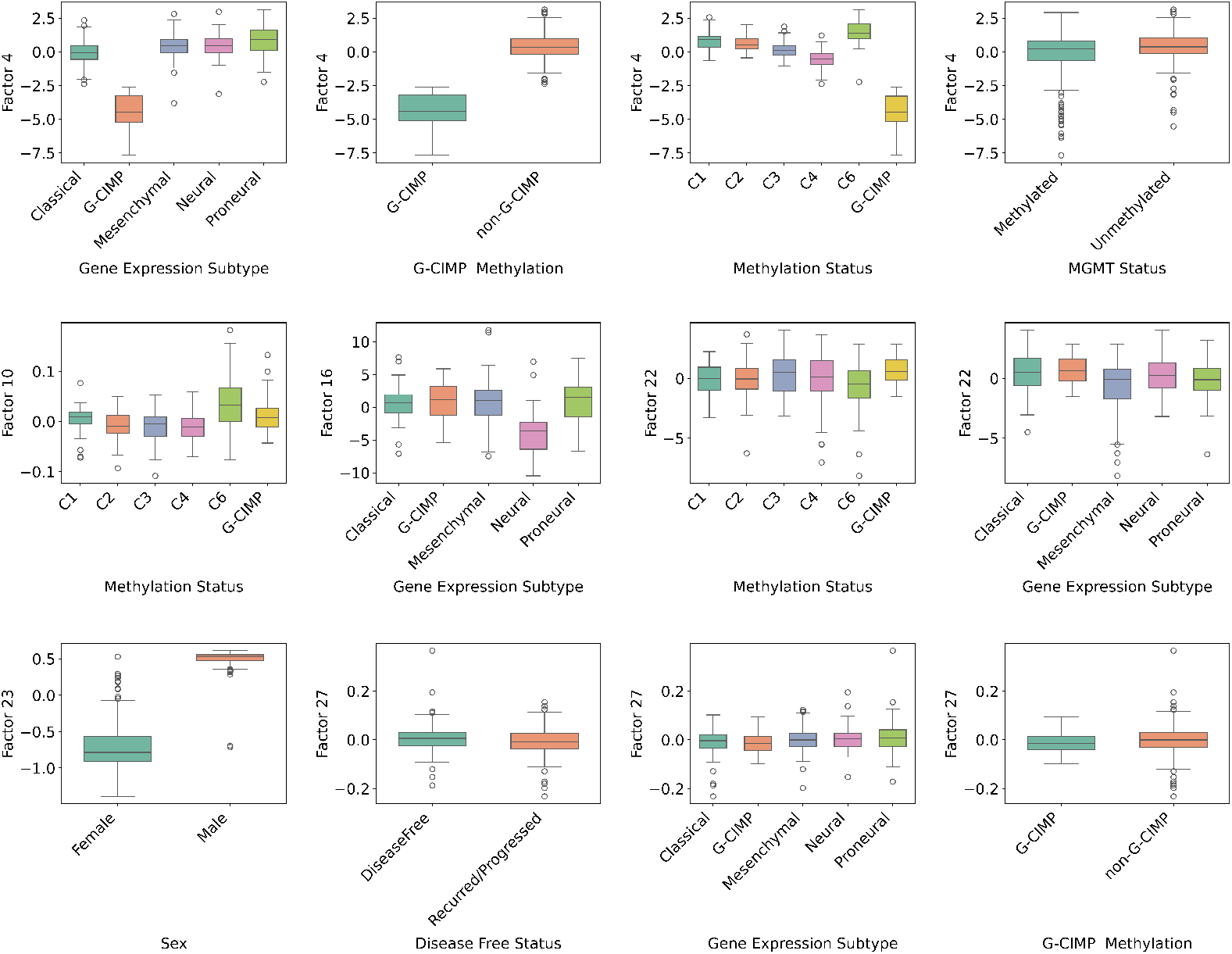
Distribution of selected latent factors across key clinical covariates in the GBM dataset. Each box represents the distribution of a latent factor within a clinical category. Differences in factor distributions highlight potential biological or clinical associations captured by CLING.

The same number of final factors *K*_final_ inferred by CLING was used to initialize other compared methods (MOFA, PCA) to ensure consistent representational capacity.

